# Alterations in resting-state network dynamics along the Alzheimer’s disease continuum: a combined MEG-PET/MR approach

**DOI:** 10.1101/2020.05.18.101683

**Authors:** D. Puttaert, N. Coquelet, V. Wens, P. Peigneux, P. Fery, A. Rovai, N. Trotta, J-C. Bier, S. Goldman, X. De Tiège

## Abstract

Human brain activity is intrinsically organized into resting-state networks (RSNs) that transiently activate or deactivate at the sub-second timescale. Few neuroimaging studies have addressed how Alzheimer’s disease (AD) affects these fast temporal brain dynamics, and how they relate to the cognitive, structural and metabolic abnormalities characterizing AD.

We aimed at closing this gap by investigating both brain structure and function using magnetoencephalography (MEG) and hybrid positron emission tomography-magnetic resonance (PET/MR) in 10 healthy elders, 10 patients with Subjective Cognitive Decline (SCD), 10 patients with amnestic Mild Cognitive Impairment (aMCI) and 10 patients with typical Alzheimer’s disease with dementia (AD). The fast activation/deactivation state dynamics of RSNs were assessed using hidden Markov modeling (HMM) of power envelope fluctuations at rest measured with MEG. HMM patterns were related to participants’ cognitive test scores, whole hippocampal grey matter volume and regional brain glucose metabolism.

The posterior default-mode network (DMN) was less often activated and for shorter durations in AD patients than matched healthy elders. No significant difference was found in patients with SCD or aMCI. The time spent by participants in the activated posterior DMN state did not correlate significantly with cognitive scores. However, it correlated positively with the whole hippocampal volume and regional glucose consumption in the right temporo-parietal junctions and dorsolateral prefrontal cortex, and negatively with glucose consumption in the cerebellum.

In AD patients, alterations of posterior DMN power activation dynamics at rest correlate with structural and neurometabolic abnormalities. These findings represent an additional electrophysiological correlate of AD-related synaptic and neural dysfunction.

## Introduction

Alzheimer’s disease (AD) is a degenerative brain disease that is the most common cause of dementia, accounting for 60 to 70% of all cases. Worldwide, about 50 million people are affected by dementia and its prevalence is expected to reach 82 million in 2030 (https://www.who.int/en/news-room/fact-sheets/detail/dementia).

Clinically, typical AD is mainly characterized by an early episodic memory impairment (i.e., difficulty to remember new facts and events) and by other cognitive difficulties such as, e.g., a dysexecutive syndrome ^1^. With the evolution of the disease, patients suffering from AD progressively develop substantial deficits in all cognitive and psychomotor functions, become bed-bound and require around the clock care ^2^.

Pathological changes begin years before symptoms onset in the AD, and usually first affect the parts of the brain involved in episodic learning (e.g., the hippocampus) and then propagate to other connected brain areas (for a review, see ^3^). They are characterized by the abnormal accumulation of two main waste proteins: the extracellular aggregates of amyloid-β peptides leading to the formation of senile plaques and the intracellular aggregates of hyperphosphorylated tau proteins leading to neurofibrillary tangles (see ^4,5^ for a descriptive of the progressive cerebral deposition). Extracellular amyloid-β deposition and senile plaques are considered to induce synaptic dysfunction and neural hyperactivity leading to network-level dysfunction, and then neuronal loss ^6^. Intracellular hyperphosphorylated tau protein and neurofibrillary tangles lead to the disruption of axonal microtubule architecture ^7^ contributing to abnormal neural and synaptic activity ^8^. Although debated (see, e.g., ^9–11^), it is hypothesized that these protein aggregates act synergistically to drive, by cell-to-cell transmission, the pathological progression of AD and neural death from the hippocampus to connected brain areas mainly encompassing the default-mode network (DMN). The DMN is composed of a set of brain regions (i.e., precuneus, posterior cingulate cortex (PCC), medial prefrontal cortex (MPFC), temporoparietal junctions (TPJs) and hippocampus) whose coordinated activity is higher when the mind is not engaged in specific behavioral tasks (i.e., during spontaneous cognitive processes at undirected rest) and lower during focused attention on the external environment or during goal-directed tasks ^12^. The brain areas that compose this network are involved in different high-level cognitive functions such as autobiographical memory, envisioning the future, theory of mind or moral decision making ^12^. Interestingly, there is a significant overlap between DMN regions and brain areas with high levels of amyloid-β and tau protein deposition, brain atrophy and hypometabolism in AD, which suggests that the DMN is particularly vulnerable to AD pathology ^13^.

The evolution towards an established dementia due to AD is progressive, and preclinical (e.g., subjective cognitive decline (SCD)) and predementia (e.g., mild cognitive impairment (MCI)) stages should be identified before clinical criteria of dementia are fully developed, knowing that not all SCD and MCI manifestations will evolve toward dementia. SCD is defined as a self-perceived decline in cognitive capacity despite normal performance at neuropsychological assessment and no significant impact on everyday activities ^14^. About 25% of people declaring SCD may develop MCI due to AD neurodegeneration, and about 10% will convert to dementia in the next 4 years ^15^. The risk of developing MCI is 4.5 times higher in people with SCD than in healthy subjects without cognitive complaint ^16^. For its part, MCI is characterized by concerns regarding a change in cognitive capabilities confirmed by an objective deficit (i.e. performance below the norms for the reference population) in one or more cognitive function(s) without an impact on everyday activities ^17^ (for a recent review, see ^18^). MCI may stem from different etiologies (e.g., degenerative as in AD, frontotemporal dementia and dementia with Lewy bodies, vascular dementia or depression), but people with amnestic MCI (aMCI, which primarily affects memory) are more likely to develop typical AD than those with non-amnestic MCI (which affects other cognitive functions) who are at higher risk of developing non-AD dementia ^19^. The estimates for MCI to AD conversion range from about 10 to 40% of patients over 5 years ^20^. These two preclinical/predementia stages thus represent a critical period during which disease-modifying treatments could slow down or even stop the degenerative progression towards AD and, consequently, dramatically improve the patients’ quality of life.

In the last decades, neuroimaging techniques such as structural magnetic resonance imaging (MRI), [^18^F]-fluorodeoxyglucose positron emission tomography (FDG-PET) and amyloid-as well as tau-PET imaging have been progressively incorporated in the list of neuroimaging markers contributing to AD management (see, e.g., ^21–24^), besides others such as cerebrospinal fluid (CSF) biomarkers. They can also contribute to the identification of people at risk of developing AD ^17,21,25^. Apart from those techniques, other neuroimaging approaches are emerging as potential interesting markers of modified brain activity in the preclinical stages. Among those, resting-state functional magnetic resonance imaging (rsfMRI), which measures the spontaneous fluctuations of blood oxygenation level-dependent (BOLD) signals in brain regions at rest (i.e., in the absence of any explicit or goal-directed task), is considered as an emerging AD-continuum marker (for reviews, see ^26,27^). Spontaneous BOLD signals typically covary between different brain areas, which is thought to reflect large-scale resting-state functional networks (i.e., the so-called resting-state networks (RSNs)) whose anatomical architecture is close to task-based functional networks ^28,29^. Among changes in RSNs uncovered by rsfMRI, evidence for AD-related alterations of DMN resting-state functional connectivity (rsFC) appears as a promising in-vivo marker of AD and its preclinical/predementia stages ^30–32^. Still, the use of fMRI along the AD continuum suffers from two important pitfalls. First, fMRI provides an indirect measure of neuronal activity via the neurovascular coupling. However, vascular changes (e.g., in vessel hemodynamics, angiogenesis, vascular cell function as well as blood-brain barrier permeability; for a review, see ^33^) are an early preclinical feature of AD pathology leading to modifications in cortical blood flow even before the onset of clinical symptoms ^34^. Thus, it cannot be ruled out that parts of rsfMRI changes observed along the AD continuum actually reflect vascular rather than neuronal changes *per se*. Second, the temporal resolution of fMRI is relatively poor (at the level of the second), which strongly impedes the characterization of the temporal and spectral dynamics of human brain activity. This means that subtle AD-related changes in neural dynamics will less likely be detected with fMRI ^35^. This explains why electrophysiological techniques such as magnetoencephalography (MEG) are increasingly recognized as potentially valuable markers in AD ^36,37^. MEG is a non-invasive neuroimaging technique measuring the extra-cranial magnetic fields produced by electrical neural activity ^38^. Contrary to fMRI, it provides a direct measure of neural activity together with an exquisite temporal resolution (of the order of milliseconds), allowing proper investigation of the temporal oscillatory dynamics of human brain activity, and is not influenced by age- and pathology-related changes in vascular coupling. MEG studies demonstrated that RSNs (including the DMN) emerge from large-scale correlation patterns in the slow fluctuations of band-limited power envelope estimated over several minutes, particularly in the alpha and beta frequency bands, confirming the neural basis of RSNs uncovered by rsfMRI ^39–45^. In addition, the high temporal resolution of MEG allows uncovering the transitory dynamics of rsFC within and across RSNs. Indeed, it has been shown that the brain spontaneously alternates between short, second-scale periods of high coupling among RSN nodes (within and across networks) and periods where only a subset of these RSN nodes interact ^46^. The DMN appears to play a central role in this dynamic functional integration of brain networks at rest ^42,47^. Importantly, these supra-second RSN dynamics could emerge from rapid transitions between recurrent brain states ^47,48^.

To the best of our knowledge, only one MEG study investigated so far the AD-related changes in the fast state dynamics underlying RSN activity ^49^, using an approach based on hidden Markov modeling (HMM). HMM identifies transient state configurations by classifying distinct patterns of power envelope (co)variance consistently repeating in time (for a review, see ^50^). From MEG data, about 6–8 transient recurring states lasting 50–200 ms are typically disclosed with a spatial topography quite similar to that of the main RSNs ^48,49,51,51^. Results obtained in AD patients demonstrated that a state of increased power in the DMN (including bilateral inferior parietal lobes, medial prefrontal cortex and lateral temporal cortices) was visited less often and for shorter periods of time in participants with AD than in matched healthy subjects ^49^. This finding suggests that spontaneous DMN activation is destabilized by AD neurodegeneration ^49^. In that study, though, the diagnosis of AD was clinical (e.g., little information was given about clinical AD criteria except for the mini-mental state examination (MMSE) and the Montreal cognitive assessment (MoCA)). Patients with preclinical/predementia stages (SCD or MCI) were also not investigated ^49^. Furthermore, the reported DMN activation state did not cover the typical posterior midline cortices (i.e., the precuneus and PCC) ^49^, possibly due to methodological issues related to source reconstruction ^53^. And finally, aberrant DMN state dynamics were not linked to cognitive alterations or established structural or functional neuroimaging markers of AD such as regional grey matter reduction ^17,21,22^ or regional brain glucose hypometabolism as measured with FDG-PET ^17,21,22^. These steps appear necessary to validate the use of MEG envelope HMM as a potential electrophysiological marker of AD.

In the present multimodal neuroimaging study, we applied HMM on MEG power envelope activity ^48^ to further characterize the alterations in RSN state dynamics along the AD continuum, and to determine their link with AD-related changes in memory function, hippocampal volume and regional brain glucose metabolism. Resting-state neuromagnetic activity was investigated in healthy elders and in patients with SCD, aMCI and typical AD. MEG power envelopes were reconstructed using Minimum Norm Estimation (MNE, see below) that allow uncovering the characteristic posterior midline cortices of the DMN ^53^. Furthermore, a hybrid PET-MR combining structural cerebral MRI and FDG-PET imaging was administered just after the MEG session. To better understand the relationships between AD pathophysiology and alterations in state dynamics, we computed correlations between altered HMM state characteristics and behavioral cognitive scores, as well as recognized markers ^21,22^ of AD such as the hippocampal volume and regional brain glucose metabolism. We expected (i) to replicate the findings of an abnormal DMN activation dynamics in AD ^49^, (ii) that reconstruction of the posterior midline cortices of the DMN would bring novel information about AD-related changes in the dynamic stability of that core human brain network, (iii) that inclusion of preclinical and predementia stages would highlight subtle modifications in the dynamics of human brain activity at rest that could serve as potential markers of the evolution toward an AD pathology, and (iv) that cross-modal correlation analyses would provide novel insights into the understanding of the pathophysiological mechanisms at the origin of these AD-related changes.

## Results

Forty participants were included in this study: 10 healthy elders (age mean and standard deviation : 70.2 ± 5.55 years; 6 females), 10 patients with SCD (71.7 ± 7; 4 females), 10 patients with aMCI (74 ± 5.47; 6 females) and 10 patients with AD (72.3 ± 7.7; 6 females) characterized by abnormal levels of amyloid-β and tau proteins in the CSF. The participant groups did not differ in age, gender ratio, years of education and handedness. Their demographic and clinical assessment data are detailed in Table 1.

**Table 1.** Participants’ demographic and clinical data. Values are presented as mean ± SD (standard deviation). Statistically significant group effects are indicated in bold. Abbreviations: HE: healthy elders, SCD: Subjective Cognitive Decline, aMCI: amnestic Mild Cognitive Impairment, AD: Alzheimer’s disease with dementia, GDS: Geriatric Depression Scale (short form), MMSE: Mini Mental State Examination, CDR: Clinical Dementia Rating scale, CSF: cerebrospinal fluid, p-tau: phosphorylated tau, sMRI: structural magnetic resonance imaging, WHV: whole hippocampal volume, FCSRT: Free and Cued Selective Reminding test, FRs: free recalls (/48), TRs: total recalls (/48), DR: delayed recall (/16), ISC: Index of Sensitivity of Cueing, na: not available. Group effects were assessed with ANOVA test for continuous variables and with χ^2^ (Pearson) for categorized indices. Normal values of *CSF Aβ:* > 381 pg/mL. Normal values of *CSF tau:* < 437 pg/mL. Normal values of *CSF p-tau*: < 61 pg/mL.

|  | HE (*n*=10) | SCD (*n*=10) | aMCI (*n*=10) | AD (*n*=10) | *p* value |
| --- | --- | --- | --- | --- | --- |
| **Age, years** | $70.2 \pm 5.55$ | $71.7 \pm 7$ | $74 \pm 5.47$ | $72.3 \pm 7.7$ | .63 |
| **Gender, females** | 6 | 4 | 6 | 6 | .77 |
| **Education, years** | $14.4 \pm 3.83$ | $16.2 \pm 5.26$ | $12.5 \pm 4.08$ | $12.9 \pm 3.17$ | .2 |
| **GDS** | $1.5 \pm 2.61$ | $4.9 \pm 3.28$ | $3.8 \pm 3.73$ | $3.66 \pm 4.18$ | .25 |
| **CDR** | $0 \pm 0$ | $0 \pm 0$ | $0.5 \pm 0$ | $0.78 \pm 0.26$ | **<.001** |
| **CSF biomarkers** |  |  |  |  |  |
| CSF Aβ, pg/mL | na | na | na | $253.8 \pm 112.1$ | - |
| CSF tau, pg/mL | na | na | na | $707 \pm 334.1$ | - |
| CSF p-tau, pg/mL | na | na | na | $92.1 \pm 31.2$ | - |
| **sMRI biomarker** |  |  |  |  |  |
| WHV, mm<sup>3</sup> | $6370.4 \pm 483.3$ | $6106.7 \pm 777.1$ | $6240.7 \pm 555.5$ | $5119.8 \pm 578.5$ | **<.001** |
| **Cognitive scores** |  |  |  |  |  |
| MMSE | $28.4 \pm 0.69$ | $28.5 \pm 0.85$ | $26.11 \pm 1.83$ | $22.2 \pm 2.48$ | **<.001** |
| Forward Digit span | $5.1 \pm 1.12$ | $5.1 \pm 0.99$ | $5.2 \pm 1.13$ | $4.4 \pm 0.84$ | .289 |
| Backward digit span | $5.8 \pm 4.19$ | $4.2 \pm 1.13$ | $4.2 \pm 1.47$ | $3 \pm 0.94$ | **.07** |
| FCSRT, *sum of FRs* | $34.5 \pm 5.2$ | $30.6 \pm 7.15$ | $15.4 \pm 3.43$ | $4 \pm 2.86$ | **<.001** |
| FCSRT, *sum of TRs* | $47.7 \pm 0.7$ | $46.2 \pm 1.68$ | $36.3 \pm 6.3$ | $13 \pm 8.83$ | **<.001** |
| FCSRT, *free DR* | $13.7 \pm 1.28$ | $11.4 \pm 2.27$ | $4.2 \pm 2.3$ | $0.6 \pm 1.07$ | **<.001** |
| FCSRT, *total DR* | $16 \pm 0$ | $15.8 \pm 0.63$ | $11.6 \pm 3.77$ | $4.2 \pm 3.19$ | **<.001** |
| FCSRT, *ISC* | $98.4 \pm 4.41$ | $90.9 \pm 8.48$ | $61.9 \pm 17.56$ | $20.8 \pm 18.02$ | **<.001** |
| Doors test (part a) | $10.3 \pm 1.49$ | $11.2 \pm 1.03$ | $7.5 \pm 2.41$ | $5.7 \pm 1.56$ | **<.001** |
| Doors test (part b) | $5.8 \pm 2.09$ | $5.6 \pm 1.95$ | $4.4 \pm 2.17$ | $2.4 \pm 1.77$ | **.002** |

All participants underwent a comprehensive neuropsychological evaluation, with, among others, evaluation of episodic/working memory and global cognition through the mini-mental state examination (MMSE, Table 1). Ongoing electrophysiological brain activity was then recorded in two consecutive 5-minute resting-state sessions ^54^ (eyes opened, fixation cross) using whole-scalp MEG. Hybrid PET-MR imaging for cerebral FDG-PET and high-resolution 3D T1-weighted structural cerebral MRI (3 Tesla) were administered just after MEG.

### Network states of MEG power activation or deactivation

We inferred RSN state dynamics from MNE-reconstructed MEG signals (across all subjects and sessions) using the HMM approach ^48^. Whole-brain wideband (4–30 Hz) source power envelopes were reduced to 8 transient recurrent states. This HMM along with the Viterbi algorithm returned a binary time series of state activation/deactivation associated with each state ^55^. The partial correlation of these time series with brain envelope signals allowed disclosing the topographical distribution of state power associated with each state ^48^. This correlation measures the degree of regional power increase/activation or decrease/deactivation during state visits. High positive (respectively negative) correlation indicated increased (respectively decreased) power envelope when the brain visited that state.

Figure 1 presents the state power maps (see *Methods* section for a description of the statistical threshold applied) of the 8 HMM transient states. Topographically speaking, State 1 involved a network configuration in which there was, upon visit, a power increase in the so-called executive network and a concomitant power decrease in the sensorimotor network. State 2 was characterized by a power decrease in bilateral angular gyri. State 3 featured increased power in the right auditory cortex and decreased power in the left lateral temporal cortex, whereas State 5 was characterized by the opposite pattern (i.e., increased power in left auditory cortex and decreased power in the right posterior temporal cortex). States 4 and 8 were marked by a power increase in the sensorimotor network, coupled with a power decrease in visual cortices for State 4. State 6 was characterized by an increase in power in brain regions corresponding to the posterior DMN (i.e., bilateral TPJs and precunei). Finally, there was increased power in visual cortices and decreased power in the ventrolateral prefrontal cortex (VLPFC) in State 7.

**Figure 1.**
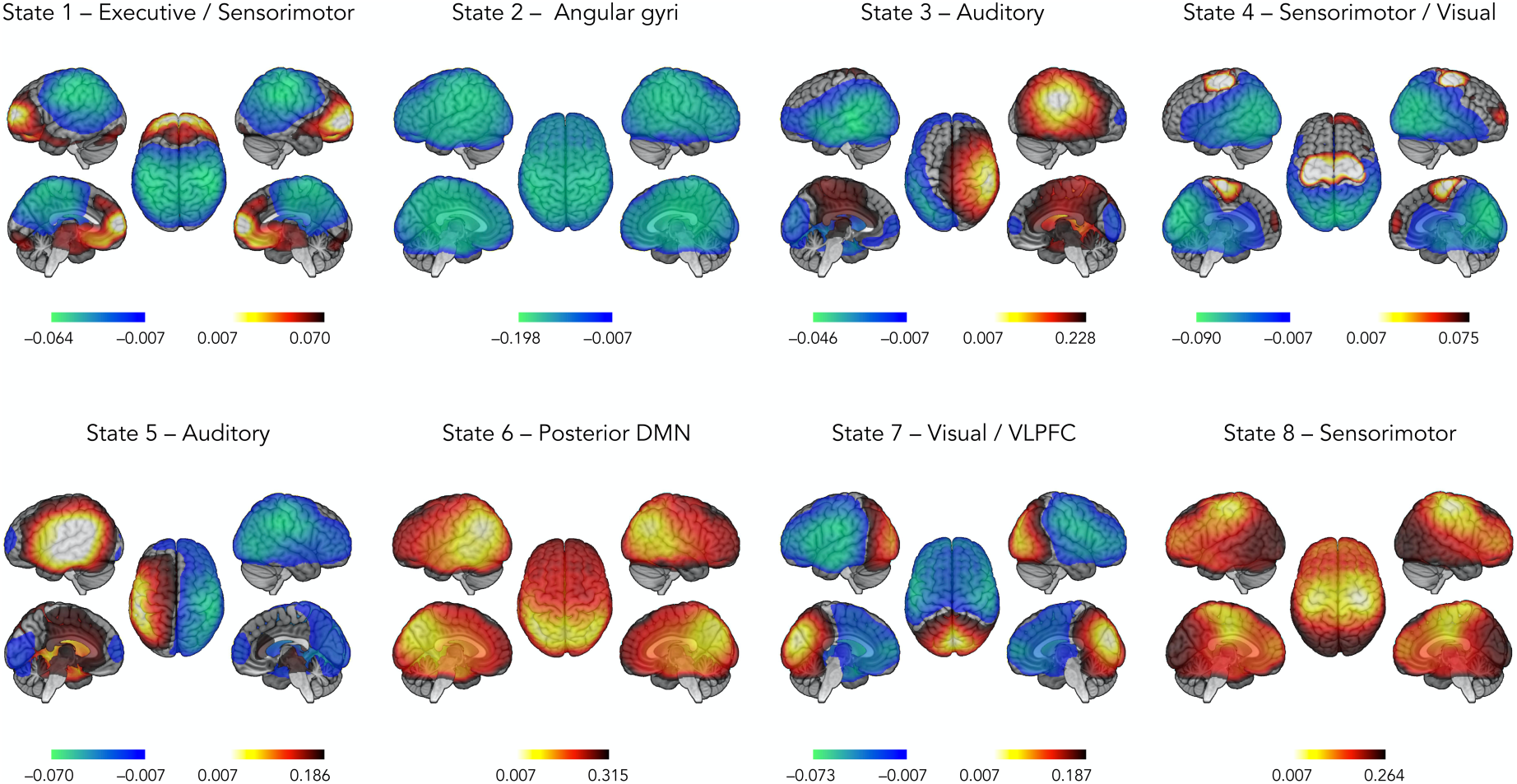
State power topography of the 8 HMM transient states. The red scale indicates positive correlation values between the envelope and the state time course (i.e., a power increase while visiting that state), whereas the blue scale indicates negative correlation values (i.e., a power decrease). Abbreviations: DMN: default-mode network, VLPFC: ventrolateral prefrontal cortex.

### Detection of altered state temporal properties

The state activation/deactivation signals also allow computing relevant state temporal parameters such as mean lifetime (MLT, i.e., the mean time spent in each state on a single visit), fractional occupancy (FO, i.e., the fraction of the total recording time that the brain spent in each state) and mean interval length (MIL, i.e., the mean time interval between two visits in the same state) ^48^. Group-level (i.e., healthy elders, SCD, aMCI and AD) differences for temporal parameters were assessed using non-parametric ANOVA (Kruskal-Wallis) with Tukey’s post-hoc tests on ranks. Significance was set at *p* < 0.05 Bonferroni controlled for the number of independent states.

Figure 2 displays the temporal parameters assessing the transience and stability of each state. Overall, MLT across the 8 states varied between 100 and 225 ms, which is in line with previous MEG envelope HMM studies ^48,49,51,51^. The only significant group effects emerged for MLT and FO of the posterior DMN activation state (State 6). The effect on MLT was explained by patients with AD spending significantly less time in that state than healthy elders (*p* < 0.001, Bonferroni corrected; healthy elders, 216.88 ms ± 54.56 ms; AD patients, 109.24 ms ± 34.85 ms); differences with SCD (176.13 ms ± 61.96 ms) and MCI (185.85 ms ± 55.46 ms) patients did not reach significance, even at *p* < 0.05 uncorrected. Likewise, FO in the posterior DMN state was significantly lower in patients with AD than in healthy elders (*p* < 0.01, Bonferroni corrected; healthy elders, 11.78 % ± 4.95 %; AD patients, 4.55 % ± 4.07 %), but not significantly lower for SCD (6.55 % ± 3.00 %) and MCI (11.28 % ± 5.06 %) patients. No significant group differences emerged for MIL at the corrected level. A tendency for higher MIL in between visits to State 6 in the AD group was observed, but only reached uncorrected significance against the MCI group probably due to large inter-subject variability among AD patients (Figure 2; healthy elders: 3.57 s ± 5.64 s, SCD: 6.15 s ± 9.75 s, MCI: 2.15 s ± 1.29 s; AD: 13.73 s ± 24.13 s). Overall, these findings suggest that AD is associated with a significant decrease in the stability of posterior DMN dynamics compared to healthy elders.

**Figure 2.**
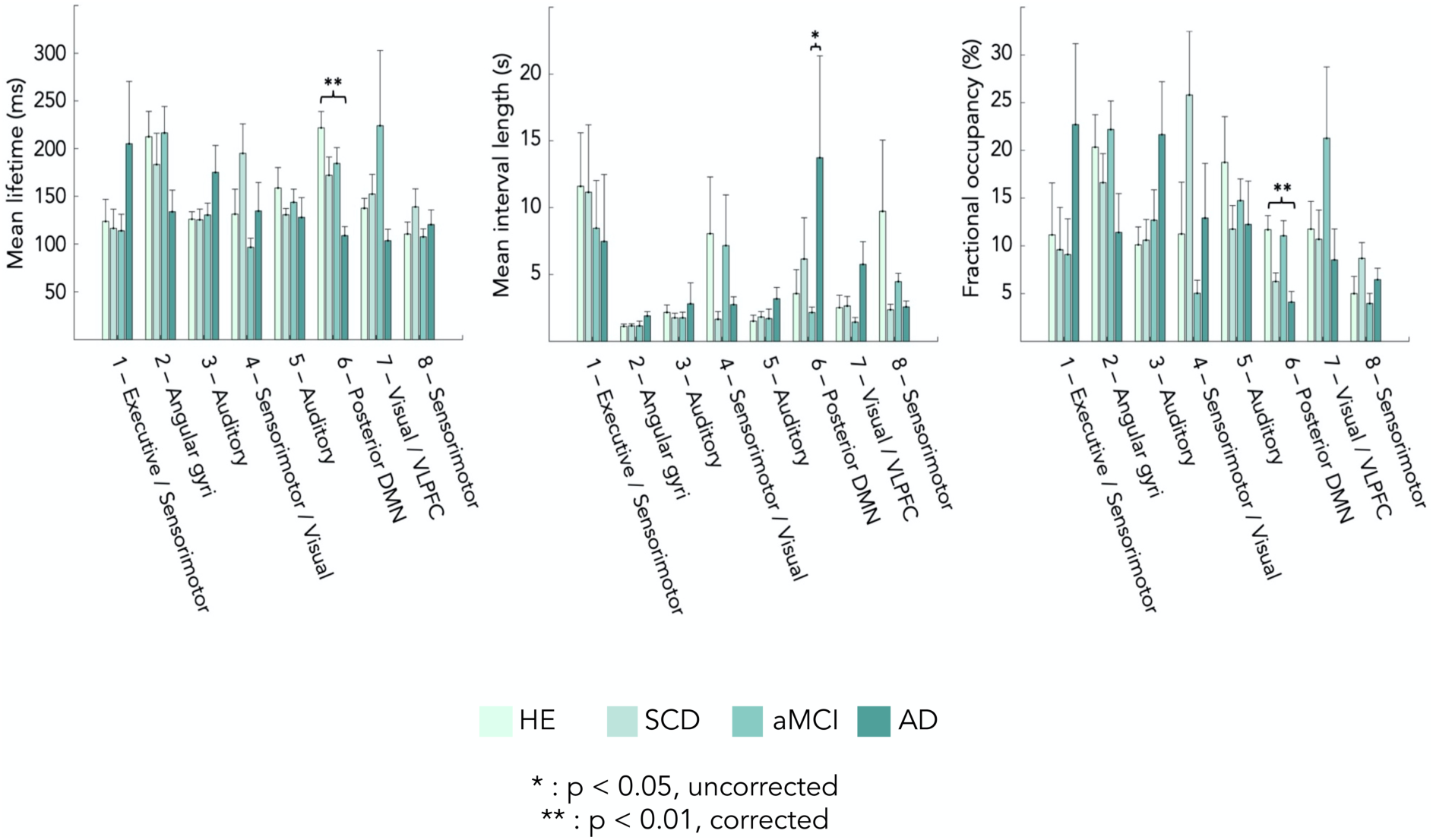
Group differences in state temporal parameters. From left to right: mean lifetime (MLT), mean interval length (MIL) and fractional occupancy (FO). Abbreviations: HE: healthy elders, SCD: Subjective Cognitive Decline, aMCI: amnestic-Mild Cognitive Impairment, AD: Alzheimer’s disease with dementia, DMN: default-mode network, VLPFC: ventrolateral prefrontal cortex. * *p* < 0.05, uncorrected; ** *p* < 0.01, Bonferroni corrected for 7 independent states. Standard error is represented above each bar.

Based on these results, we selected MLT and FO of the posterior DMN activation State 6 for further correlation analyses with (i) cognitive test scores (Table 1), (ii) the whole hippocampal volume (Table 1), and (iii) regional brain glucose metabolism.

### Correlation between State 6 temporal parameters and cognitive test scores

No significant (all *p*s > 0.05, corrected) correlation (computed across the 40 participants) was observed between State 6 temporal parameters (MLT and FO) and cognitive test scores (episodic and visual memory, working memory and MMSE). Thus, no conclusive link could be drawn between the temporal instability of posterior DMN activation and cognitive impairment.

### Correlation between State 6 temporal parameters and hippocampal volume

The left and right hippocampal grey matter volumes were identified by tissue segmentation of participants’ structural MRI using FreeSurfer v6.0 (Martinos Center for Biomedical Imaging, Massachusetts, USA). A significant positive correlation (r = 0.314; *p* = 0.048) was observed between MLT of State 6 and the whole hippocampal volume (i.e., summed over left and right volumes). Indeed, the bigger the whole hippocampal volume was, the longer was the time spent in State 6 (Figure 3). No significant correlation was observed with the FO of State 6 (r = 0.18, *p* = 0.26).

**Figure 3.**
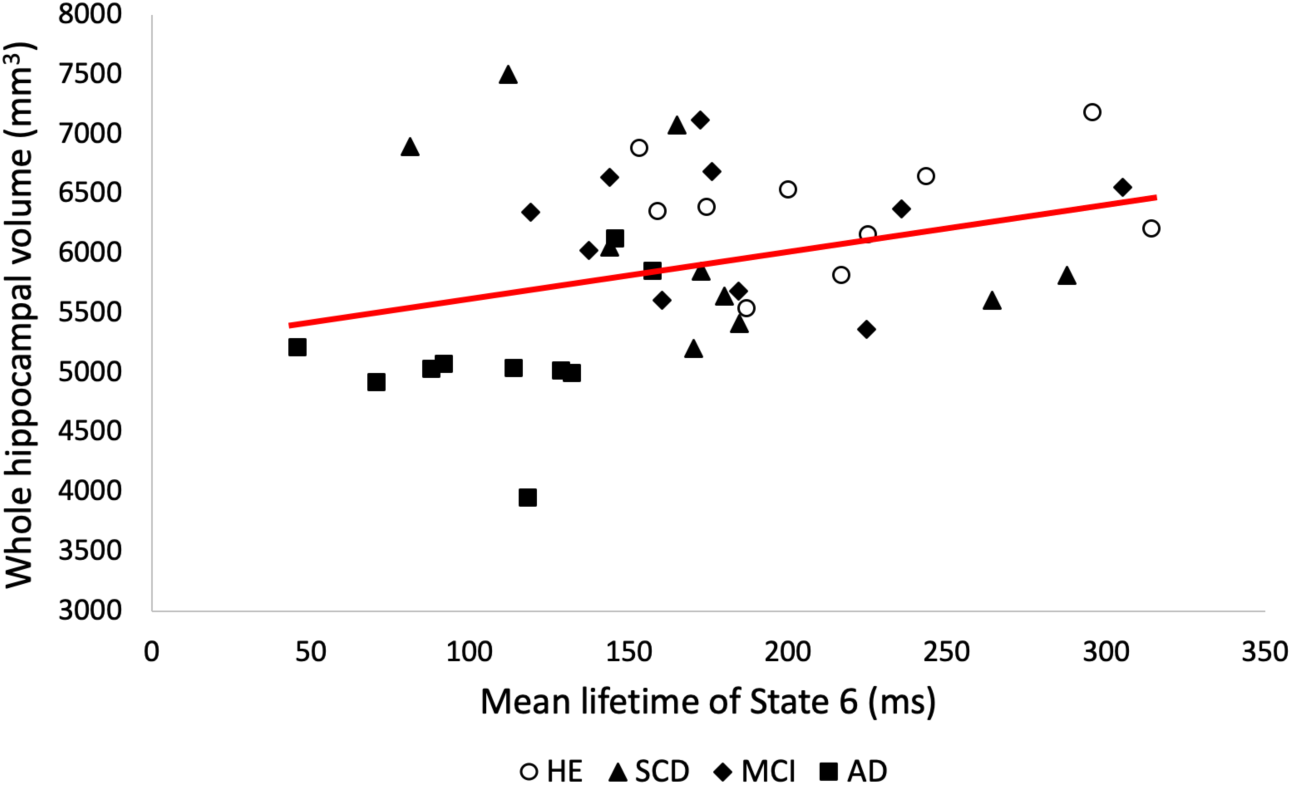
Correlation between mean lifetime of State 6 and whole hippocampal volume. Abbreviations: HE: healthy elders, SCD: Subjective Cognitive Decline, aMCI: amnestic-Mild Cognitive Impairment, AD: Alzheimer’s disease with dementia.

### Correlation between State 6 temporal parameters and regional cerebral glucose metabolism

Statistical parametric correlation maps of State 6 temporal parameters and regional brain glucose metabolism measured with FDG-PET were obtained using SPM12 (Wellcome Centre for Neuroimaging, London, UK). We identified positive correlations with MLT in the right dorsolateral prefrontal cortex (DLPFC), right angular gyrus, right supramarginal gyrus and right superior temporal gyrus (voxel-based correlation *T* tests at *p* < 0.001 uncorrected, with cluster threshold at *k* > 100 voxels ^56^; see Figure 4, top and Table 2). This means that the less time the participants spent activating their posterior DMN, the lower was the glucose consumption in a right fronto-parietal network. A significant negative correlation was also observed with MLT in bilateral cerebellum (*p* < 0.001 uncorrected, *k* > 100 voxels; see Figure 4, bottom and Table 3). A similar result held for FO, which was negatively correlated with regional brain glucose metabolism in the right cerebellum (Lobule V) (*p* < 0.001 uncorrected, *k* > 100 voxels; see Figure 5 and Table 4). Therefore, the less time the participants spent activating their posterior DMN, the higher was the glucose metabolism in these cerebellar areas.

**Table 2.** Brain areas showing positive correlation between mean lifetime of State 6 and regional brain glucose metabolism. We report the MNI coordinates of the local cluster peaks in Figure 4 alongside the corresponding *T* value.

| *Anatomical regions* | *MNI coordinates (mm)* |  |  | *T value* |
| --- | --- | --- | --- | --- |
|  | *x* | *y* | *z* |  |
| DLPFC | 36 | 20 | 32 | 5.07 |
| Right angular gyrus | 46 | -56 | 34 | 4.1 |
| Supramarginal gyrus | 60 | -62 | 32 | 4.06 |
| Right superior temporal gyrus | 54 | -44 | 22 | 3.86 |

**Table 3.**
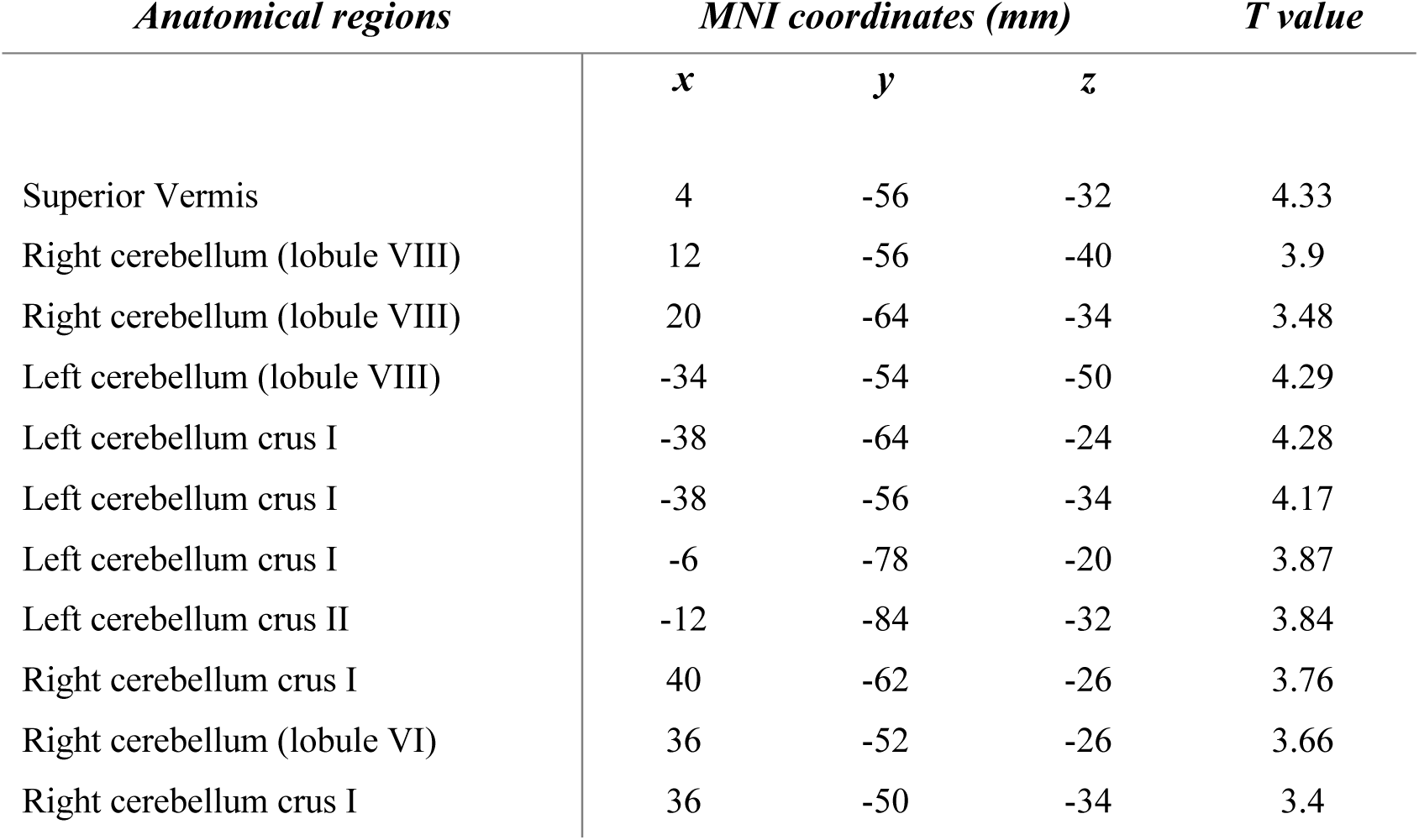
Brain areas showing negative correlation between mean lifetime of State 6 and regional brain glucose metabolism in all participants. We report the MNI coordinates of the local cluster peaks in Figure 4 alongside the corresponding *T* value.

| *Anatomical regions* | *MNI coordinates (mm)* |  |  | *T value* |
| --- | --- | --- | --- | --- |
|  | *x* | *y* | *z* |  |
| Superior Vermis | 4 | -56 | -32 | 4.33 |
| Right cerebellum (lobule VIII) | 12 | -56 | -40 | 3.9 |
| Right cerebellum (lobule VIII) | 20 | -64 | -34 | 3.48 |
| Left cerebellum (lobule VIII) | -34 | -54 | -50 | 4.29 |
| Left cerebellum crus I | -38 | -64 | -24 | 4.28 |
| Left cerebellum crus I | -38 | -56 | -34 | 4.17 |
| Left cerebellum crus I | -6 | -78 | -20 | 3.87 |
| Left cerebellum crus II | -12 | -84 | -32 | 3.84 |
| Right cerebellum crus I | 40 | -62 | -26 | 3.76 |
| Right cerebellum (lobule VI) | 36 | -52 | -26 | 3.66 |
| Right cerebellum crus I | 36 | -50 | -34 | 3.4 |

**Table 4.** Brain areas showing negative correlation between fractional occupancy of State 6 and regional brain glucose metabolism in all participants. We report the MNI coordinates of the local cluster peaks in Figure 5 alongside the corresponding *T* value.

| *Anatomical regions* | *MNI coordinates (mm)* |  |  | *T value* |
| --- | --- | --- | --- | --- |
|  | *x* | *y* | *z* |  |
| Right cerebellum (lobule V) | 36 | -50 | -22 | 5.23 |

**Figure 4.**
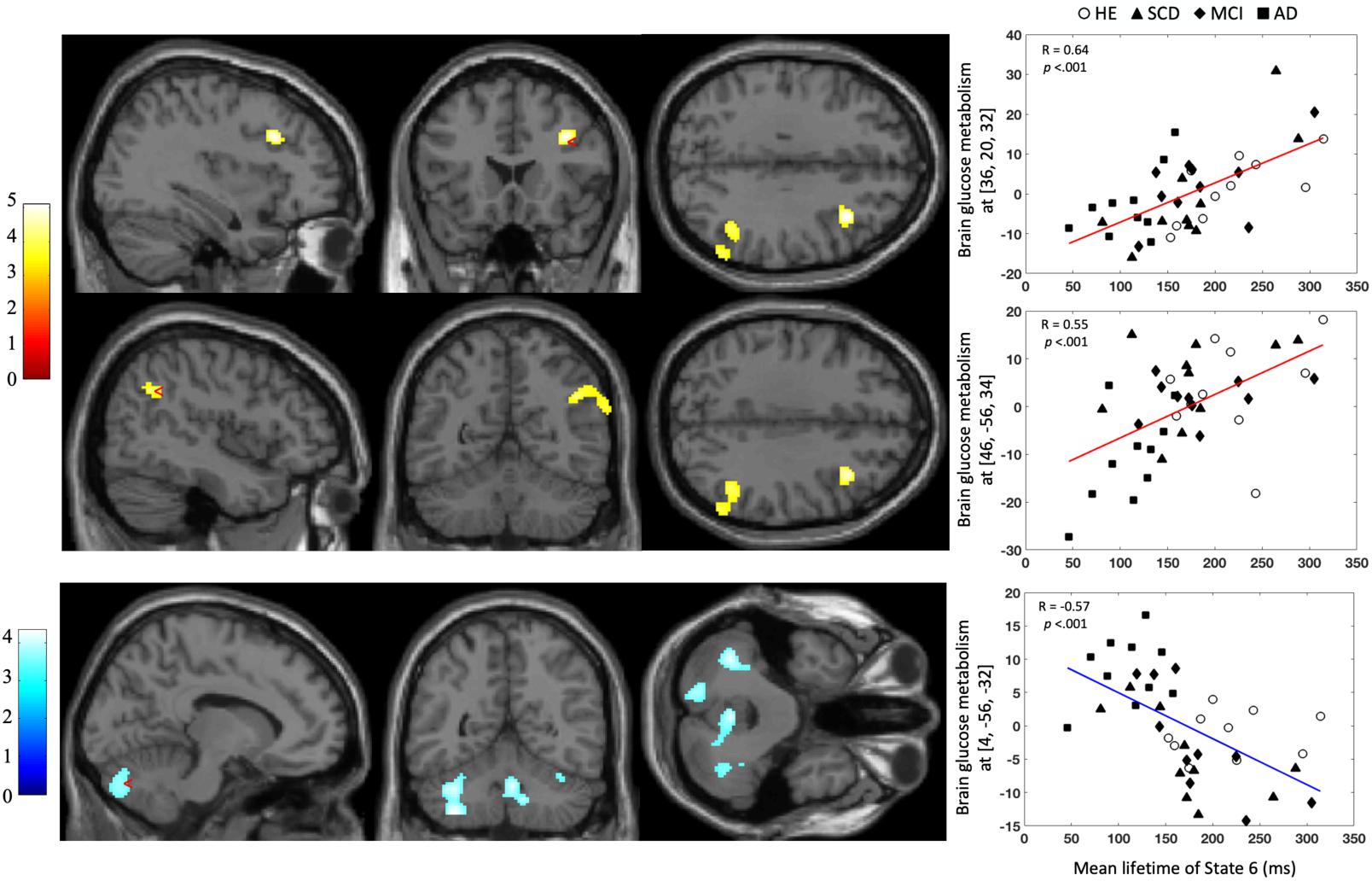
Statistical parametric *T* maps assessing significant positive (Top) and negative (Bottom) correlations between mean lifetime of State 6 and FDG-PET signals across the 40 participants. Images are thresholded at *p* < 0.001 uncorrected with a cluster threshold at 100 voxels.

**Figure 5.**
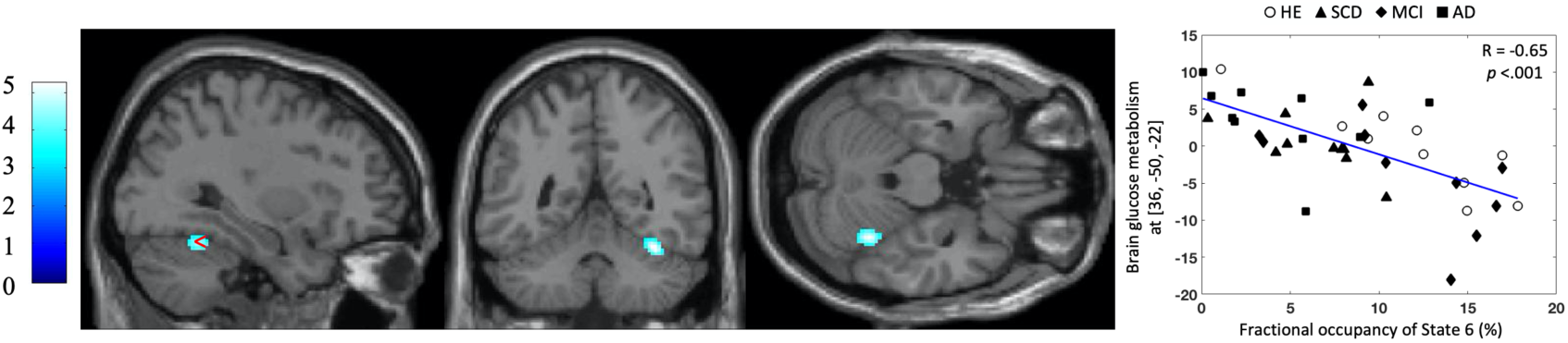
Statistical parametric *T* maps assessing significant negative correlation between fractional occupancy of State 6 and FDG-PET signals across the 40 participants. Images are thresholded at *p* < 0.001 uncorrected with a cluster threshold at 100 voxels.

## Discussion

This study evidenced less frequent visits and less time spent in a transient recurrent state of activated posterior DMN in patients with AD than in matched healthy elders. No difference in transient resting-state brain dynamics was found between patients in pre-clinical (i.e., SCD) or pre-dementia (i.e., MCI) stages and either the patients with AD or the healthy elders. The time spent by the participants in the activated posterior DMN state was positively correlated with the whole hippocampal volume and with glucose consumption in right TPJ and DLPFC, was negatively correlated with glucose consumption in the cerebellum, but did not correlate with memory and global cognition scores.

The 8 transient recurrent HMM states disclosed in our participants exhibited spatial and temporal patterns rather similar to those previously described ^48,49,51,52,57^. Importantly, we also found one state (State 6) characterized by activated bilateral TPJs and precunei when visited, corresponding to an activated posterior DMN state. Involvement of posterior midline cortices such as the precuneus was not disclosed in previous MEG HMM studies that identified similar DMN states ^48,49^. In particular, it was not observed in the DMN state reported in the MEG envelope HMM study of AD ^48^. This is presumably related to the use of MNE rather than beamforming for source reconstruction in this study, as it was shown that the former is better suited to image midline posterior cortices in functional integration studies ^53^.

Patients with AD showed a dynamical destabilization of this posterior DMN activation characterizing State 6. There was a significant reduction in MLT and FO of that state in patients with AD compared with matched healthy elders (along with a trend for higher MIL in AD patients). These finding are in line with those of a previous MEG HMM study performed in AD patients ^49^, except that we further identified altered dynamics in the bilateral precunei. Furthermore, we investigated here a population of AD patients with established abnormal amyloid and tau protein levels in the CSF. Our results therefore reinforce the hypothesis that, compared to healthy elders, AD is characterized by a significant reduction in the temporal stability of power activity within the posterior DMN. Considering the pathognomonic decrease in (static) resting-state functional connectivity within the DMN typically found in AD patients (for a review, see, e.g., ^58^), this finding suggests established AD-related dementia is accompanied by a reduction in both DMN integration and dynamic activation.

The time spent by participants in the activated posterior DMN state did not correlate with memory and global cognition scores (episodic and visual memory, working memory and MMSE scores). This finding suggests that the aberrant AD-related destabilization of resting-state posterior DMN activation may not be directly linked to cognitive impairment. This lack of conclusive correlation with the cognitive profile of AD patients was also observed in a rsfMRI study of dynamic functional integration ^59^. A specific hypothesis for this is further developed below.

By contrast, significant correlations were found with the whole hippocampal grey matter volume and regional brain glucose metabolism. A positive correlation was indeed observed with the whole hippocampal grey matter volume, demonstrating that destabilization of the activated posterior DMN state was associated with hippocampal atrophy. Hippocampal atrophy, as evidenced by structural cerebral MRI, is one of the best validated and widely used (but still imperfect) AD markers (for reviews, see, e.g., ^60–62^). In this disease, the decrease in hippocampal volume is strongly associated with hippocampal neuronal loss and number of neurofibrillary tangles caused by intracellular aggregates of hyperphosphorylated tau proteins (see, e.g., ^63,64^). Also, studies showed that early in the symptomatic disease process, AD patients exhibit reduced (static) functional integration between hippocampi and DMN posterior midline cortices ^58^. Our correlation between hippocampal structure and DMN dynamics therefore supports the hypothesis that AD-related changes in transient resting-state brain dynamics are intrinsically linked with AD pathophysiology. The alteration in posterior DMN resting-state stability may therefore be considered as an additional electrophysiological correlate of the AD-related synaptic and neural dysfunction. A positive correlation was also observed between the time spent by participants in the activated posterior DMN state and glucose consumption in right TPJs and DLPFC, demonstrating that the higher the dynamical instability for that state, the less was the glucose metabolism in these brain areas associated with a right fronto-parietal network. A first hypothesis to explain this finding is that the typical AD-related decrease in regional glucose metabolism would be asymmetric (right-sided) in our population of AD patients. However, this is not supported by subtractive voxel-based SPM analyses comparing FDG-PET data of healthy subjects with those of AD patients (data not shown for conciseness, methods used similar to ^65–67^). A second hypothesis to consider is that this right-hemispheric lateralization was due to the right-handedness of the majority of our participants. In this view, the AD-related reduction in DMN activation would be associated with a reduction of glucose metabolism in the ventral attention network (VAN) ^68^, which tends to be right-lateralized in right-handed people. This would be reminiscent of a previously described AD-related reduction in the intrinsic anticorrelation between the DMN and the VAN ^69–72^. Finally, a negative correlation was found with glucose consumption in the cerebellum (bilateral cerebellar hemispheres and superior vermis), showing that the more destabilized the activated posterior DMN state, the higher was the cerebellar glucose metabolism. The role of cerebellum in AD is not fully understood, but this result may be in line with some studies suggesting its implication in compensatory mechanisms of AD pathology (for a review, see ^73^). For example, better memory performance has been correlated with an increase in cerebellar activity during encoding in AD patients ^74^.

To our knowledge, this study is the first to investigate the dynamic properties of resting brain activity using an HMM approach applied to MEG power envelope signals from potentially preclinical and predementia stages of AD (i.e., in SCD and aMCI patients). However, analyses failed to identify any significant difference in HMM state temporal parameters between these groups and healthy elders or AD patients. A possible explanation is that the dynamic electrophysiological abnormalities in the DMN are too subtle to be detectable in preclinical and prodromal stages of AD, and would rather progress sharply at the latest stage. As a consequence, our relatively small sample size could induce a lack of significant sensitivity. We cannot also exclude the hypothesis that our samples of patients with SCD and aMCI were mostly composed of the percentage of patients who will not evolve toward AD. Indeed, the evolution of preclinical and predementia stages is extremely variable between individuals (see, e.g., ^75^). MCI and SCD stages are heterogeneous conditions that may be caused by different pathologies (see, e.g., ^76–79^), so it can be expected that the majority of SCD and MCI patients included in our study will never progress to AD. Another explanation for these negative results may thus be that these two groups of participants are on average more alike to healthy subjects, possibly overshadowing a continuous progression of DMN alteration from SCD to AD. A longitudinal follow-up of our participants will be essential to investigate the potential evolution of these preclinical/predementia groups to AD, and to compare the initial profile of transient resting-state brain dynamics between individuals who remained stable or evolved to AD after a few years.

A key issue associated with our findings is to determine whether the AD-related changes in transient resting-state brain dynamics are linked to the possible effects of pathological aging on intrinsic (i.e., task-independent neural processes) functional brain integration or on spontaneous cognitive processes (i.e., task-dependent neural processes), or on their complex interactions. Indeed, whether spontaneous brain activity and functional integration reflect intrinsic (i.e., task-independent) neural processes (e.g., maintenance of homeostasis or the integrity of anatomical connections) or extrinsic (i.e., task-dependent) neural processes, or both, remains an open question (for a review, see, e.g., ^80^). As certain subtypes of spontaneous cognitive processes are detectable in time-varying functional connectivity measurements ^80^, it could be hypothesized that AD-related changes in transient resting-state DMN activation dynamics observed in this study might pertain to the emerging literature about changes in the occurrence of mind-wandering episodes and in the content/type of spontaneous cognitive processes observed in pathological aging ^81–83^. Elders indeed have a tendency to experience less mind-wandering episodes and with different cognitive contents than young adults ^84–87^, with a further decline in AD patients ^81^. AD-related alterations in spontaneous cognitive processes seem indeed highly plausible considering the key role of the DMN in some spontaneous cognitive processes (mind wandering, etc.) and the peculiar DMN vulnerability (amyloid-β and tau protein deposition, atrophy, hypometabolism) in AD (for a review, see ^88^). Based on those considerations, the destabilization of posterior DMN activations at rest might represent an electrophysiological correlate of the decrease in mind-wandering episodes reported in AD. They might therefore reflect more AD-related changes in extrinsic rather than intrinsic brain dynamics. The absence of link between rather stable (i.e., across hours, days) cognitive deficits ^89,90^ and transient (a few hundred ms) resting-state posterior DMN dynamics might bring additional support to this hypothesis. Further studies combining behavioral and electrophysiological investigations of transient brain processes along the AD continuum are therefore mandatory to clarify this intrinsic/extrinsic issue.

The present study suffers from several limitations. First, the sample size of each group (healthy elders, SCD, aMCI and AD) was rather modest. Still, we demonstrated similar HMM results than the previous HMM MEG study performed in AD ^49^, using the same number of AD patients but with established abnormal amyloid and tau protein CSF levels. This replication therefore suggests that the observed differences in transient resting-state DMN dynamics between AD patients and healthy elders are robust. Still, as discussed above, this modest sample size may have impacted our sensitivity to detect possible subtle differences between pre-clinical/pre-dementia stages and healthy elders or AD patients. Further studies including large numbers of participants along the AD continuum are therefore mandatory to confirm our findings. Second, as mentioned before, SCD and MCI stages are heterogeneous conditions characterized by different possible progressions to AD or non-AD dementia, stabilization or even normalization of their cognitive profile ^75^. That could partially explain the lack of significant results for these preclinical and predementia stages. Indeed, our study is cross-sectional and, consequently, the accurate percentage of SCD and MCI patients who will develop AD in few years is unknown. Future longitudinal investigations are needed to explore the validity of these results. Third, a refinement of the criteria defining these preclinical and predementia stages, notably with the measure of CSF biomarkers (i.e., amyloid-beta, total-tau and phosphorylated-tau), could be particularly useful in that context. Indeed, a previous study showed that CSF biomarker score provided a good estimate of the risk of MCI to AD dementia conversion up to 6 years in comparison with lower CSF biomarker score ^91^. Additionally, an AD-positive metabolic pattern, identified with FDG-PET, in MCI was previously considered as the best predictor of conversion to AD (in comparison with typical CSF and MRI markers). A combined approach could be more efficient to clarify the definition of SCD, MCI as well as the risk of progression to AD dementia, as previously demonstrated ^92^. Such voxel-based comparison of each patient’s FDG-PET data with that of those of the healthy elders’ group was not performed in the present study due to (i) the low number of healthy elders and the need to have a large group of healthy subjects (>50) to increase the sensitivity/specificity of such highly multivariate FDG-PET analyses ^93^, and (ii) because of the high-resolution and MRI-based correction for attenuation of our FDG-PET data, which complicate the use of FDG-PET data available in large database (e.g., ADNI). Furthermore, a previous study has shown that healthy elders at risk for AD (i.e., with Aβ-positive defined by using C-Pittsburgh compound B PET), contrary to healthy elders with Aβ-negative, presented an increase in rsFC between the precuneus and the bilateral inferior parietal lobules in theta and delta frequency bands ^94^. This finding suggests an association between these electrophysiological rsFC changes and local cerebral amyloid deposition ^94^. Further research should investigate the dynamic resting-state changes in healthy elders at risk of developing AD in order to clarify the relation with AD pathology. Finally, recent works have shown that the use of cryogenic MEG systems induced an underestimation of the level of frontal functional integration due to inhomogeneities in the MEG sensor-brain distance ^80,95^. We cannot therefore totally exclude that some AD-related changes have been underestimated in those brain regions due to a lower signal to noise ratio. The use of on-scalp MEG based on optically pumped magnetometers, which have been demonstrated to be usable for neuromagnetic investigations in humans, should therefore be privileged in the future ^96^.

In summary, this study demonstrated that abnormalities in fast brain network activation dynamics can be identified in AD using HMM analysis of resting-state MEG power envelopes. Results suggested that AD-related alterations of posterior DMN activity may be considered as an additional electrophysiological correlate of the AD-related synaptic and neural dysfunction. Whether these alterations reflect AD-related changes in intrinsic (i.e., task-independent) vs. extrinsic (i.e., task-dependent) brain activity remains an open key question.

## Materials and Methods

### Participants

Ten healthy elders, 10 SCD *plus* patients (see *Diagnostic criteria* below), 10 aMCI and 10 typical AD patients were included in this study. All participants gave their written informed consent prior to their participation in this study conducted in agreement with the Ethics Committee of CUB Hôpital Erasme (P2017/427, Brussels, Belgium).

General exclusion criteria were previous other neurological or psychiatric disorders, a modified Hachinski Ischemic Score ≥ 4 (HIS) ^97^, chronic use of psychotropic drugs or alcohol, insufficient level in French language, age younger than 55 or older than 90 years, and severe visual or hearing impairment. All individuals had a least 6 years of education and the majority of them were right-handed (except for one healthy subject who was left-handed, and one healthy subject and one aMCI who were ambidextrous) according to the Edinburgh handedness inventory ^98^.

#### Clinical evaluation

All participants were initially screened for exclusion criteria. Then, the clinical evaluation included the minimal mental scale examination (MMSE) ^99^, the short form of the geriatric depression scale (GDS) ^100^, the neuropsychiatric inventory (NPI) ^101^, the Pittsburgh sleep quality index (PSQI) ^102^ and a questionnaire assessing the quality of participants’ previous night of sleep (adapted from ^103^). An exhaustive anamnesis and a hetero anamnesis were conducted with all patients and their caregivers in order to learn more about their medical and psychiatric history. Furthermore, the clinical dementia rating (CDR) ^104^, and the basic and instrumental activities of daily living (BADLs and IADLs) ^105^ questionnaires were used to evaluate the severity of symptoms and their potential impact on everyday life. Each participant also underwent a comprehensive neuropsychological assessment comprising the evaluation of (i) verbal episodic memory with the free and cued selective reminding test (FCSRT) ^106,107^, (ii) visual episodic memory using the Doors and People test (only the Doors part was administered) ^108^, (iii) working memory by the forward and backward digit span test (Wechsler Memory Scale, WMS-III), (iv) language abilities with the ExaDé naming test ^109^, (v) executive functions and, more specifically, the flexibility sub-component using the phonemic/letter and categorical fluency test ^110^, as well as the trail making test (part B) ^111^ and the inhibition sub-component with the interference part of the Stroop color and word test ^112^, (vi) visuo-constructive abilities with the Rey-Osterrieth figure copy ^113^ and finally, (vii) speed processing and visual attention with the trail making test (part A) ^111^.

#### Diagnostic criteria

Healthy elders had a normal neuropsychological performance and no subjective memory complaint. SCD patients were defined following Jessen’s criteria including : (1) self-experienced persistent decline in cognitive capacity in comparison with a previously normal status and unrelated to an acute event, (2) normal age, gender and education-adjusted performance on standardized neuropsychological tests, (3) not to be MCI or AD as per the criteria described below, and (4) subjective cognitive decline cannot be explained by a psychiatric or neurologic disease, medical disorder or substance use ^14^. SCD *plus* were defined with subjective decline in memory rather than other domains of cognition, the onset of SCD within the last 5 years, the age at onset of SCD ≥ 60 years, concerns and feelings of worse performance than others of the same age group (evaluated here with the GDS-short form question : *“Do you feel you have more problems with your memory than most”* ^100^). The aMCI subjects were classified according to the core clinical criteria developed by the National Institute on Aging – Alzheimer’s Association (NIA-AA) workgroups ^17^ : (1) concerns regarding a change in cognition, (2) an impairment in one or more cognitive domains, (3) preservation of independence in functional abilities, and (4) to be not demented. Furthermore, according to their main memory impairment, they were all classified with the amnestic profile of MCI ^114^. Finally, the new criteria of NIA-AA ^24^ were used to define AD patients with dementia following the amyloid deposition, pathologic tau, and neurodegeneration [AT(N)] classification that groups different biomarkers by the pathologic process. Typical profile of AD patients was characterized by greater deficit of verbal episodic memory than other cognitive functions. Indeed, one study showed that 79.6% patients with typical AD cognitive profile have an autopsy-confirmed diagnosis compared to only 20.4 % for atypical AD group ^115^.

### Multimodal data acquisition

Neuromagnetic data were acquired (two sessions of 5 minutes, eyes open, sitting position, fixation cross, online band-pass filter : 0.1-330 Hz, sampling rate : 1000 Hz) by using a 306 channel whole-scalp MEG system (Triux™, MEGIN, Helsinki, Finland) placed inside a light-weight magnetically shielded room (Maxshield™, MEGIN, Helsinki, Finland; see ^116^ for more details) at the CUB Hôpital Erasme (Brussels, Belgium). The position of participants’ head was recorded continuously inside the MEG helmet by using four head tracking coils. Coils’ location and approximatively 300 head points were determined following the anatomical fiducials with an electromagnetic tracker (Fastrak, Polhemus, Colchester, Vermont, USA). Furthermore, eye movements or blinks and cardiac artifacts were captured with bipolar electrodes.

T1-weighted MRI (Repetition Time/Echo Time/Flip Angle: 8.3 ms/3.1 ms/12°, TI: 450 ms, field of view (FOV): 24 cm x 24 cm, matrix: 240 × 240, resolution: 1 mm x 1 mm x 1 mm) and FDG-PET data acquisitions were performed simultaneously on a hybrid 3 T SIGNA™ PET-MR scanner (GE Healthcare, Milwaukee, Wisconsin, USA) at the Service of Nuclear Medicine (CUB Hôpital Erasme, Brussels, Belgium). All participants fasted for at least 4 hours, were awake in an eye-closed rest and received an intravenous bolus injection of 2–5 mCi (74– 185 MBq) of FDG before PET-MR data acquisition. The time interval between FDG injection and the start of data acquisition was 40 min and the total scan duration was 20 min. PET data were reconstructed using the fully 3D iterative reconstruction algorithm VUE Point FX-S, which takes into account the time-of-flight information and the correction for the point spread function of the system. The algorithm was configured with 10 iterations, 28 subsets and a standard Z-axis filter cut-off at 4 mm. The photons’ attenuation specific of the PET acquisition was corrected with a MRI-based map (MRAC) acquired simultaneously. PET images were displayed in a 256 × 256 x 89 matrix format, with a slice thickness of 2.78 mm. The reconstructed files were downloaded in their original format (DICOM, ECAT, Interfile) for meta-information and converted in NIfTI format to be finally used for analysis.

### Data preprocessing

MEG data were filtered offline using temporal signal space separation to remove external interferences and nearby sources of magnetic artifact, but also compensate for head movements ^117^. Cardiac, ocular and system artifacts were visually identified and eliminated using an independent component analysis (FastICA algorithm with dimension reduction to 30 components, hyperbolic tangent nonlinearity function) ^118^ of the filtered data (off-line band-pass filter: 0.1–45 Hz).

The MEG forward model was computed on the basis of participants’ 3D T1-weighted cerebral MRI, which was anatomically segmented beforehand using the FreeSurfer software (version 6.0; Martinos Center for Biomedical Imaging, Massachusetts, USA). Of note, this segmentation procedure also provided the right and left whole hippocampal grey matter volumes used for further correlation analyses. FreeSurfer is classically used to obtain hippocampal grey matter volumes in the field of AD ^119–121^. The co-registration between MEG and MRI data was then performed using the 3 fiducial points (nasion and auricular points) for first estimation and the head-surface points to manually refine the surface co-registration (Mrilab, Elekta Oy, Helsinki, Finland). Then, a volumetric and regular 5-mm source grid was built using the Montreal Neurological Institute (MNI) template and non-linearly deformed onto each participants’ MRI with the Statistical Parametric Mapping Software (SPM 12, Wellcome Centre for Neuroimaging, London, UK). The three-dimensional MEG forward model associated with this source space was computed using a one-layer Boundary Element Method as implemented in the MNE-C suite.

The preprocessing of FDG-PET images was performed with SPM8 (www.fil.ion.ucl.ac.uk/spm) and spatially normalized using a specific FDG-PET aging and dementia template ^122^. Data were then smoothed with a FWHM 8-mm Gaussian kernel.

### MEG source reconstruction

A band-specific MNE ^123^ was applied to perform inverse modeling that was restricted to the 204 planar gradiometers. Five minutes of artifact-free noise measurement, obtained from an empty-room MEG recording filtered spatially with signal space separation ^117^ and spectrally in the relevant frequency range, was recorded for the estimation of the noise covariance matrix. The regularization parameter was calculated with the consistency condition (as derived in ^124^). The depth bias was corrected using a noise normalization scheme, i.e., dynamic Statistical Parametric Mapping ^123^. Each estimated three-dimensional source was projected onto its direction of maximum variance, filtered in the 4–30 Hz band, and Hilbert transformed to extract their wide-band envelope signal fed to the HMM analysis.

### Hidden Markov Modeling

We thoroughly followed the pipeline described in ^48,125^ implemented in GLEAN (https://github.com/OHBA-analysis/GLEAN), except for the use of MNE as inverse model rather than a Beamformer. The number of transient states (*K*) was set to 8 for consistency with previous MEG envelope HMM studies ^48,125^. The 8-state HMM was inferred from the wide-band (4-30 Hz) source envelope signals. These envelopes were first downsampled at 10 Hz using a moving-window average with 75% overlap (100 ms wide windows, sliding every 25 ms), leading to an effective downsampling at 40 Hz. Envelope data were demeaned and normalized by the global variance and then concatenated temporally across participants to design a group-level HMM analysis. Group-concatenated envelopes were finally pre-whitened and reduced to 40 principal components. The HMM *per se* was then applied to determine dynamic states of envelope covariance, along with the Viterbi algorithm to obtain binary time series of state activation/deactivation under the constraint of temporal exclusivity ^55^. These binary signals allowed to determine individual estimates of several state temporal parameters such as the MLT (i.e., mean duration of time intervals of active state), the FO (i.e., total fraction of time during which the state is active) and the MIL (i.e., mean duration of time intervals of inactive state).

Non-parametric ANOVA (Kruskal-Wallis) was used to detect group effects (healthy elders, SCD, aMCI, AD) on state-specific temporal properties. Significance was set to *p*<0.05 Bonferroni corrected for the number *K*-1=7 of independent states (one state being dependent upon the others due to the temporal exclusivity constraint). Tukey’s post-hoc testing on data ranks allowed to identify group differences explaining significant ANOVA.

State power maps were built as the partial correlation between HMM state activation/deactivation time series and group-concatenated source envelope signals. These correlations assessed state-specific power changes upon state activation. These maps were thresholded statistically by applying a two-tailed parametric correlation test at *p* <0.05 against the null hypothesis that Fisher-transformed correlations following a Gaussian with mean zero and standard deviation 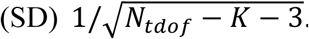. The number *N*_*tdof*_ of temporal degrees of freedom was calculated as one-quarter of the total number of time samples to take into account the overlap of 75% in the envelope downsampling. The critical *p* value was Bonferroni corrected with number of independent HMM states (i.e., 7) and the number of spatial degrees of freedom in MNE maps (estimated from the rank of the MEG forward model ^124^).

### Correlation between temporal properties of HMM states and cognition, whole hippocampal volume, and regional brain metabolism

Correlation analyses with altered state temporal parameters (i.e., disclosing a significant group effect) aimed at obtaining novel insights into the pathophysiology of their alterations in the context of AD.

First, the Pearson correlation coefficient was estimated (across the 40 participants) between altered state temporal parameters and cognitive scores. More specifically, the scores included in this analysis were : verbal episodic memory as assessed using Delayed Recall after 20 minutes, sum of free recalls, sum of total recalls and the index of sensitivity of cueing ([“sum of the 3 total recalls” – “sum of the 3 free recalls”]/[48 – “sum of the 3 free recalls”], defined by ^126^) from ^107^, the visual episodic memory with the doors and people test (only the doors part was administered) ^108^, working memory with forward and backward digit span tests (Wechsler Memory Scale, WMS-III) and finally, global cognition defined by MMSE score ^99^. The results were considered significant at *p* < 0.05 Bonferroni corrected for multiple comparisons following the number of cognitive scores considered.

A similar Pearson correlation analysis was performed with the whole hippocampal volume, computed as the sum of left and right whole hippocampal grey matter volumes obtained from structural MRI segmentation using FreeSurfer (see above). Our focus on the hippocampus was driven by that hippocampal atrophy is a well-recognized feature of AD and an established neuroimaging marker ^17,25^.

Finally, a mass-univariate version of Pearson correlation analysis was used to explore the relationship between state temporal parameters and regional cerebral glucose metabolism. We used SPM12 (http://www.fil.ion.ucl.ac.uk/spm/, Wellcome Trust Centre for Neuroimaging, London, UK) to construct two (i.e., one form MIL and one for FO) voxel-based general linear model (GLM) of the preprocessed FDG-PET data of all participants (40 scans), with the mean-centered state temporal parameter (i.e., MIL or FO) as covariate of interest. Proportional scaling was applied beforehand to remove inter-subject variation in global brain metabolism and an explicit FDG mask was used to restrict the analyses inside the brain. Separate *T* contrast analyses then searched, throughout the brain, for regions showing significant positive or negative correlations. As such, this analysis is an adaptation of the previously developed psycho-, physio-, or patho-physiological interaction analyses ^66,127^ to a specific experimental factor, i.e., state temporal parameters. Results were considered significant at *p* < 0.001 uncorrected with a cluster threshold at 100 voxels, considering the limited number of participants.

## Data availability

The datasets analyzed during this current study are available from the corresponding author upon reasonable request.

## Acknowledgments

Delphine Puttaert is a researcher of the Fonds Erasme (Brussels, Belgium). Xavier De Tiège is Postdoctoral Clinical Master Specialist at the Fonds de la Recherche Scientifique (FRS-FNRS, Brussels, Belgium). The PET-MR project at the CUB Hôpital Erasme and Université libre de Bruxelles is financially supported by the Association Vinçotte Nuclear (AVN, Brussels, Belgium). The MEG project at the CUB Hôpital Erasme is financially supported by the Fonds Erasme (Brussels, Belgium) via the research convention “Les Voies du Savoir”.

## Author contributions

D.P., S.G., P.P., J-C.B. and X.D.T. designed the study; D.P. acquired data; D.P., N.C., V.W., A.R., N.T., S.G. and X.D.T. analyzed and interpreted data; D.P., N.C., V.W., S.G. and X.D.T. wrote the manuscript, D.P., N.C., V.W., P.P., J-C.B., S.G., A.R., P.F., N.T. and X.D.T. drafted the manuscript for intellectual content, X.D.T. obtained funding.

## Additional Information

The authors declare that they have no competing financial interests.

## Notes

### Competing Interest Statement

The authors have declared no competing interest.

